# A hotspot for conformational heterogeneity driven by proline isomerisation in the androgen receptor disordered N-terminal domain

**DOI:** 10.64898/2026.07.29.741599

**Authors:** Mathilde Roth, Hélène Launay, Eva Erdmann, Veronique Receveur-Brechot, Jocelyn Ceraline, Bruno Kieffer, Célia Deville

## Abstract

The androgen receptor is a hormone-dependent transcription factor that regulates a wide range of physiological processes and plays a pivotal role in the development of prostate cancer. Its 555-residue, intrinsically disordered, N-terminal domain is involved in the modulation of transcriptional activity by recruiting co-regulators and mediating the formation of biomolecular condensates. This study reports on the characterisation of a conserved domain located in the C-terminal region of the androgen receptor N-terminal domain, at atomic level, using nuclear magnetic resonance spectroscopy. This proline rich region exhibits extensive conformational heterogeneity driven by highly populated cis proline conformers that are stabilised through interactions with adjacent aromatic residues. We demonstrate that the cis-proline population is modulated by phosphorylation as well as cancer-associated mutations. This suggests that proline driven conformational heterogeneity at the C-terminal region of androgen receptor N-terminal domain is involved in the regulatory function of this transcription factor.

## Introduction

Intrinsically disordered regions (IDRs) are highly prevalent in eukaryotic transcription factors (Wright and Dyson, 2015; Udupa et al., 2024) with over 80% of proteins in this family containing disordered segments of 30 or more amino acids (Liu et al., 2006). Among these, the nuclear receptors superfamily shares a common architecture (Figure 1). A ligand-binding domain (LBD) is responsible for hormone-dependent activation called activation function 2 (AF-2). A DNA-binding domain (DBD) recognizes hexanucleotide motifs from nuclear receptor response elements and a flexible hinge region (H) connects these structured domains. Upstream of the DBD lies the N-terminal domain (NTD), an intrinsically disordered region of variable length that holds the ligand-independent transcriptional activity called activation function 1 (AF-1) (Tora et al., 1989; Weikum et al., 2018). The androgen receptor (AR) possesses a particularly long NTD, consisting of 555 amino acids representing 60% of the full-length protein (Lavery and Mcewan, 2005). Early functional studies have firmly established the NTD as a critical component of the AR regulation: its full deletion abolishes AR activity, and several large regions including the Tau-1 (residues 102-371) and Tau-5 (361-537) transactivation domains were shown to be especially important for AR activation in a gene reporter assay (Jenster et al., 1995). Besides, LBD-truncated AR variants, which emerge in castration-resistant prostate cancer (CRPC), retain constitutive hormone-independent transcriptional activity (Céraline et al., 2004; Dehm et al., 2008; Marcias et al., 2010), though with distinct target gene programs compared to the full-length receptor (Erdmann et al., 2022). Beyond its role in AR activation, the NTD serves as a platform to integrate context-dependent regulatory signals.

**Figure 1.**
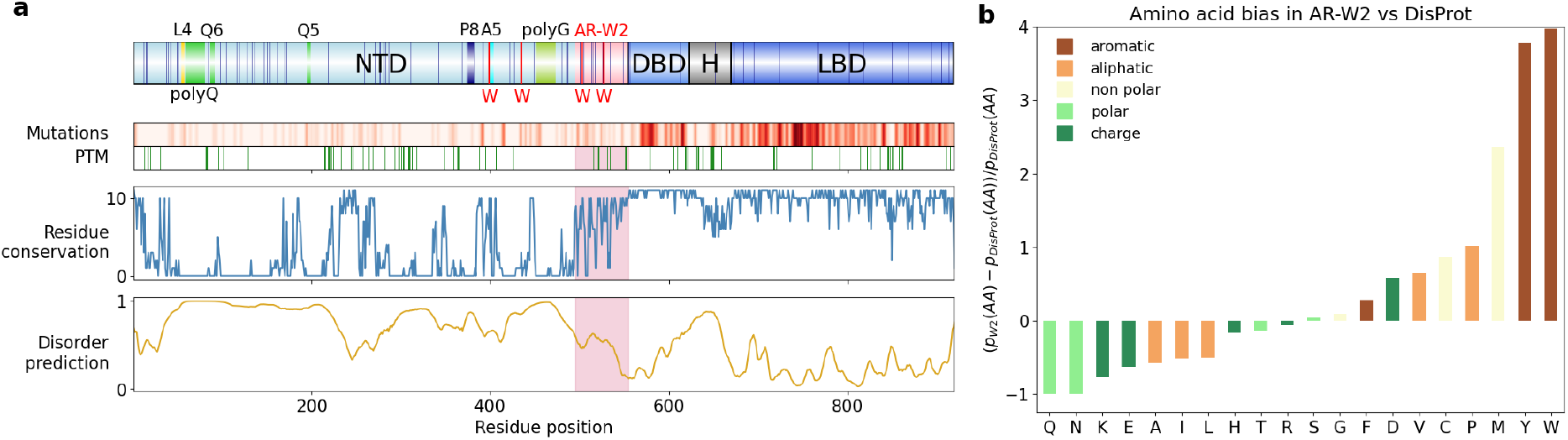
Architecture of the androgen receptor. **a.** Domain architecture of the androgen receptor. NTD: N-terminal Domain; DBD: DNA Binding Domain: H: Hinge; LBD: Ligand Binding Domain. Homopolymers over 4 residues are shown as coloured bars and prolines as blue lines. AR-W2 region is highlighted in lightpink. Position of individual post-translational modification (PTM) sites are shown as green bars. **b**. Amino acid bias in AR-W2 compared to the DisProt database of intrinsically disordered regions.

The NTD dynamically modulates AR activity through multiple layers of regulation according to a complex IDP molecular grammar (Holehouse and Kragelund, 2024) where sequence polymorphism and post translational modifications plays a pivotal role. AR-NTD harbors two long homopolymeric tracts, a polyglutamine and a polyglycine, both of 23 amino acids in the reference P10275 Uniprot sequence and whose lengths vary across individuals. Critically, shorter polyglutamine tracts correlate with enhanced AR transcriptional activity, while expansions beyond 40 amino acids are linked to diminished transactivation and the neurodegenerative disorder spinal bulbar muscular atrophy (Kennedy’s disease) (Meszaros et al., 2022). At the post-translational level, the NTD hosts half of AR’s 83 post translational modification sites (PTM) that affect receptor stability, localisation and transcriptional activity (Gioeli and Paschal, 2012), enabling rapid adaptation to cellular signals, as documented in the PhosphoSitePlus database. Furthermore, although most of AR mutations associated to prostate cancer and androgen insensitivity syndrome (AIS) are located in folded domains, affecting DNA or ligand binding properties, approximately one-quarter missense mutations are found in AR-NTD (Figure 1).

Biophysical and functional studies have been undertaken to elucidate how AR-NTD intrinsic disorder encodes for regulatory versatility and decipher the molecular grammar of this region. NMR spectroscopy revealed, local transient secondary structures as well as interactions with coregulators and small-molecule inhibitors (De Mol et al., 2016; De Mol et al., 2018; Eftekharzadeh et al., 2019) for fragments ranging from the N-terminus to the beginning of the polyglycine (residue 448). Moreover, AR-NTD drives higher-order AR assembly: aromatic residues promote liquid-liquid phase separation (LLPS) (Basu et al., 2023; Xie et al., 2022), while the polyglutamine tract and conserved KELCKAVSVSM motif mediate aggregation and reversible amyloid fibril formation (Asencio-Hernández et al., 2014; Oppong et al., 2017).

Sequence alignment across vertebrates (from Homo sapiens to cartilaginous fish) reveals that while most of AR-NTD shows low conservation, as expected for an IDR, a few highly conserved regions stand out, notably a long 60 amino acid stretch directly adjacent to the DBD. This conserved C-terminal region of AR-NTD (residues 495-555), hereafter termed AR-W2, displays unique sequence features distinguishing it from typical IDRs. Notably, two tryptophane residues, usually the least abundant residue in IDRs, are perfectly conserved within AR-W2 region: W503 and W527 (Figure S1). AR-W2 is enriched in prolines, aromatic (especially in tyrosines and tryptophans) but also aliphatic and non polar residues, an atypical feature when comparing to the average IDR composition from Disprot (Figure 1.b).

AR-W2 is particularly rich in prolines, distributed across the sequence and not clustered in proline rich regions. Usually described as a disorder-promoting residues because it lacks the amide hydrogen necessary to form stabilising hydrogen bonds in *α*-helices or *β* -sheets, prolines are particularly abundant in intrinsically disordered regions (Theillet et al., 2013). Due to their cyclic nature, their cis conformer of the peptidyl proline peptide bond is more favoured than for other amino acids, with a baseline population of 5-10%, both in folded and disordered regions. Besides increased population of cis isomer, the cis/trans interconversion kinetics spans a wide range of time scales from seconds to hours. This slow inter-conversion effectively partitions the conformational ensemble into distinct subpopulations, each with different structural and possibly functional properties. Exchange between trans and cis proline conformation can be accelerated by prolyl isomerases, which reduce the timescale to the millisecond range and such proline isomerases are known to be coregulators of the androgen receptor (Periyasamy et al., 2007; Yong et al., 2007; Periyasamy et al., 2010; La Montagna et al., 2012) although so far only Pin1 was precisely mapped to interact with AR-NTD (La Montagna et al., 2012; Leung et al., 2021).

In addition to its sequence peculiarities, AR-W2 displays a high level of polymorphism with 17 mutations identified in prostate cancer and AIS as well as four post-translational modification sites: three phosphorylation sites at S515, Y535 and Y552 and a sumoylation site at K521 (Figure S1), all described as modulators of androgen receptor activity (Guo et al., 2006; Dai et al., 2010; Gioeli and Paschal, 2012). Comparison with the C-terminal region of the NTD of other steroid receptors shows no sequence conservation and different amino acid composition (Figure S2) indicating AR-specific function for this region.

AR-W2 was previously shown to reduce the affinity of androgen receptor DNA-binding domain (DBD) for androgen response elements *in vitro* (Liu et al., 2003), likely due to its net negative charge that generates repulsive electrostatic interactions with DNA, a common feature of IDRs adjacent to DNA binding domains, where such interactions are proposed to prevent non specific DNA binding (Udupa et al., 2024). AR-W2 was also shown to decrease AR transactivation *in cellulo* (Liu et al., 2003) and to be involved efficient recruitment of p160 coactivator NCOA2 (Need et al., 2009) but the underlying molecular mechanisms remain unknown. The description of AR-W2’s conformational landscape thus represents a first step to link AR-W2 sequence features to its regulatory function. NMR spectroscopy was used to characterise conformational heterogeneity in AR-W2 at atomic resolution (Jensen et al., 2014). We identified hotspots for cis proline conformations and in particular Gly-Pro-Aro motifs that, compared to the symmetrical Aro-Pro-Gly motifs, display remarkably high cis Gly-Pro peptide bond populations. This Gly-Pro-Aro cis proline preference is associated to restricted aromatic side chain dynamics. Finally, we observe that the Gly-Pro cis conformer population is modulated by post-translational modifications and cancer associated mutations.

## 1. Methods

### 1.1 Sequence analysis

322 androgen receptor sequences were retrieved from the SwissProt (12 sequences) and RefSeq databases (including 17 NP curated sequences and 294 XP computationally predicted sequences). Protein sequences were aligned using MAFFT (Katoh et al., 2019) and manually curated with Jalview (Waterhouse et al., 2009). Because of the presence of homopolymers of glutamines and glycines of variable length, different versions of AR numbering co-exist in the literature. Here we use AR uniprot entry P10275 as a reference for residue numbering.

AR missense mutations were retrieved from the LOVD database (version 3000-30b, downloaded on September 22nd 2025). Local mutation density was calculated using a sliding 5 residues window, taking into account occurence of mutations in the database.

### 1.2 Peptide synthesis and preparation

The following peptides were chemically synthesised:

P5 A**P**DVWY**P**GGMVSRV**P**Y**P**S**P**TC 499-519

P2 CVKSEMG**P**WMDSYSG**P**YGD 519-537

P2inv CVKSEM**WGP**MDSYSG**P**YGD 519-537

P2pY535 CVKSEMG**P**WMDSYSG**PpY**GD 519-537

P2G525D CVKSEM**DP**WMDSYSG**P**YGD 519-537

where pY is a phosphorylated tyrosine.

Peptide Synthesis was performed on a 433A Peptide Synthesizer (ABI) using Fmoc chemistry. Crude peptides were purified by reverse phase HPLC on a kinetex 5*µ*m EVO C18 100Å 250 × 21.2 mm column from Phenomenex. Purified peptides were controlled by Mass Spectrometry (XEVO G2-XS Qtof Waters).

Lyophilised peptides were dissolved to a concentration of 2-6mM in NMR buffer (50mM phosphate buffer pH=6.5, 150mM NaCl, 2-10mM TCEP). The pH of the solution was ajusted to 6.5 with NaOH.

The absence of effect of residual TFA in the peptide solutions was assessed by removing TFA using 3 repeated cycles of peptide lyophilization after addition of a 0.1 M HCl solution (Andrushchenko et al., 2007). Decrease of TFA concentration was quantified from 1D ^19^F spectra and no significant changes were observed on ^1^H-^15^N and ^1^H-^13^C HSQC spectra.

### 1.3 Protein expression and purification

AR-W2 and AR-W2DBD fragments were produced fused to a protein tag composed of a N-terminal 6xHis tag, a maltose binding protein (MBP) and a Tobacco Etch Virus (TEV) cleavage site. Plasmids were transformed into *Escherichia coli* BL21 (DE3) cells. For production of unlabelled or isotopically labelled protein for NMR studies, cells were grown in LB medium or M9-minimal media (containing 1g/L ^15^NH_4_Cl and 1g/L of ^13^C glucose for ^15^N and ^13^C labelling respectively). Protein expression was induced by addition of 0.5mM IPTG when the culture reached an OD_600_ of 0.6-0.8 and expression was carried out overnight at 20°C. Cell were harvested by centrifugation and pellets were stored at -20°C until purification.

Pellets were resuspended in lysis buffer (50mM Tris-HCl pH=7.5, 150mM NaCl, 10% glycerol, 1mM DTT and EDTA-free cOmplete protease inhibitor cocktail from Roche). Cells were lysed by sonication and the lysate was clarified by ultra-centrifugation (125 000 g, 1 hour, 4°C). The supernatant was injected on a 5mL HisTrap FF crude column (GE Healthcare) equilibrated with buffer A (50mM Tris-HCl pH=7.5, 150mM NaCl, 15mM imidazole, 1mM DTT). Protein was eluted with buffer B (50mM Tris-HCl pH=7.5, 150mM NaCl, 200mM imidazole, 1mM DTT). The His-MBP fusion was cleaved by the TEV protease and the sample was dialysed against 15mM imidazole buffer to remove imidazole excess. The protein cleaved tag and His-tagged TEV protease were captured with Ni-NTA resin and the AR-W2 protein was further purified by size exclusion chromatography on a Superdex 75 HiLoad 16/600 column equilibrated with GF buffer (50mM phosphate pH=7.0, 150mM NaCl). Fractions containing the protein of interest were pooled and immediately complemented with 1mM TCEP and protease inhibitors (EDTA-free cOmplete from Roche). Protein was concentrated by cycles of centrifugation on 3kDa cut-off concentrators. Purification steps and protein purity were assessed via SDS-PAGE and mass spectrometry.

### 1.4 NMR spectroscopy

All NMR samples were prepared using 8-10% D_2_O, 50mM phosphate buffer (pH=6.5), 150mM NaCl, 1-10mM TCEP. NMR spectra were acquired on Bruker AVANCE 700 and 950MHz (^1^H) spectrometers equipped with cryogenic probes at a temperature of 278K to reduce amide proton exchange with water. DSS (sodium 2,2-dimethyl-2-silapentane-5-sulfonate) was used to reference chemical shifts (*δ*(^1^*H*)_*DSS*_ = 0). Spectra were processed in Topspin and analysed using the CcpNmr software suite(Skinner et al., 2016). Backbone assignment of AR-W2 was performed using ^1^H-^15^N-HSQC, HNCA, HN(CO)CA, HNCACB, HN(CO)CACB, HNCO, HN(CA)CO experiments. For peptides, ^1^H-^15^N and ^1^H-^13^C-HSQC were recorded at natural abundance of ^15^N and ^13^C using peptide concentration between 2 and 6mM for total experiment times ranging from 4 to 16 h for ^1^H-^13^C-HSQC and 10 to 30 h for ^1^H-^15^N-HSQC. Assignments were partly transferred from the ^1^H-^15^N-HSQC of AR-W2 and subsequently confirmed and completed using NOESY (200 ms mixing time), TOCSY (90 ms mixing time) and ^1^H-^1^3C-HSQC-TOCSY (60 ms mixing time) experiments. Proline conformation for each of the identified states was determined by examining the proline C_*β*_ and C_*γ*_ chemical shifts difference which is in the range of 5 and 10 ppm for trans and cis prolines respectively (Schubert et al., 2002). Peak volumes were estimated by fitting correlation peaks with a lorentzian line shape in CCPNMR. Grouped fitting was performed for massifs and peaks for which overlap prevented accurate intensity determination were manually excluded from the analysis. Population of each cis proline conformer was estimated by calculating the volume ratios of correlation peaks corresponding to the same nuclei in distinct conformations, both for the proline residue itself and for all neighbouring residues for which proline isomerisation resulted in distinct correlation peaks. The uncertainty in the population estimates was calculated as the standard deviation of these values, corrected using the appropriate Student’s t coefficient for a two-sided 95% confidence interval. Uncertainty contributions arising from the signal-to-noise ratio of individual peaks were found to be negligible relative to the variability between population estimates obtained from different correlations and were therefore neglected.

## 2. Results

### 2.1 AR-W2 is intrinsically disordered in absence and presence of the adjacent DBD

AR-W2 was studied either in isolation or in the context of AR-DBD by purifying two fragments: AR-W2 (residues S495–P555) and AR-W2DBD (residues S495–G628). The intrinsically disordered nature of AR-W2 was confirmed by the low chemical shift dispersion of proton chemical shifts on ^1^H-^15^N-HSQC spectra. The ^1^H-^15^N-HSQC spectrum of AR-W2DBD confirmed the disordered nature of AR-W2 in presence of this adjacent DBD (Figure S3). Besides, the presence, for AR-W2 and AR-W2DBD spectra, of 4 distinct peaks in the tryptophans HN*ε* region for these fragments containing only 2 tryptophans suggest conformational heterogeneity. Additional evidence was provided by the presence of numerous minor correlation peaks (up to 30% intensity of major peaks) in the ^1^H-^15^N-HSQC NMR spectrum. Mass spectrometry confirmed sample integrity, excluding proteolysis and leading us to hypothesize that these minor states arise from proline isomerization.

A standard set of 3D NMR experiments enabled complete assignment of the major conformation of AR-W2 where all prolines adopt trans conformation, as evidenced by their C_*β*_ chemical shifts. AR-W2 contains 9 prolines distributed throughout its sequence, and partial assignment of minor peaks confirmed their localization near proline residues. However, unambiguous and complete assignment was hindered by low populations of some cis states falling below 3D NMR detection limits and spectral overlap, particularly in ^13^C dimensions where poor chemical shift dispersion hindered unambiguous assignment. To systematically characterize these minor conformational states, we employed a reductionist approach using peptide fragments (Smet et al., 2004) spanning AR-W2’s proline-rich regions.

### 2.2 Evenly distributed prolines induce large conformational heterogeneity in AR-W2

We synthesised 2 peptides comprising 7 of the 9 prolines of AR-W2: P5(Ala499-Cys519) contains P500, P505, P513, P515 and P517 and P2 (Cys519-Asp537) contains P526 and P534 (Figure 2a). ^1^H-^15^N and ^1^H-^13^C HSQC spectra were recorded at natural abundance of carbon and nitrogen. ^1^H-^15^N HSQC spectra of P5/P2 overlapped closely with AR-W2 (except terminal residues), validating their use as minimal models (Figure 2b, Figure S4). Proton NOESY, TOCSY and ^1^H-^13^C TOCSY HSQC spectra allowed to validate assignment of the major conformation and to assign all minor conformations. This complete assignment of minor conformations in P5/P2 confirmed 7 distinct cis proline conformations (Figure 2a-b), with variable segmental effects from long-range propagation (12 residues affected) for P526/P534 to very localized effects (2–3 residues) for P513/P517 (Figure 2c).

**Figure 2.**
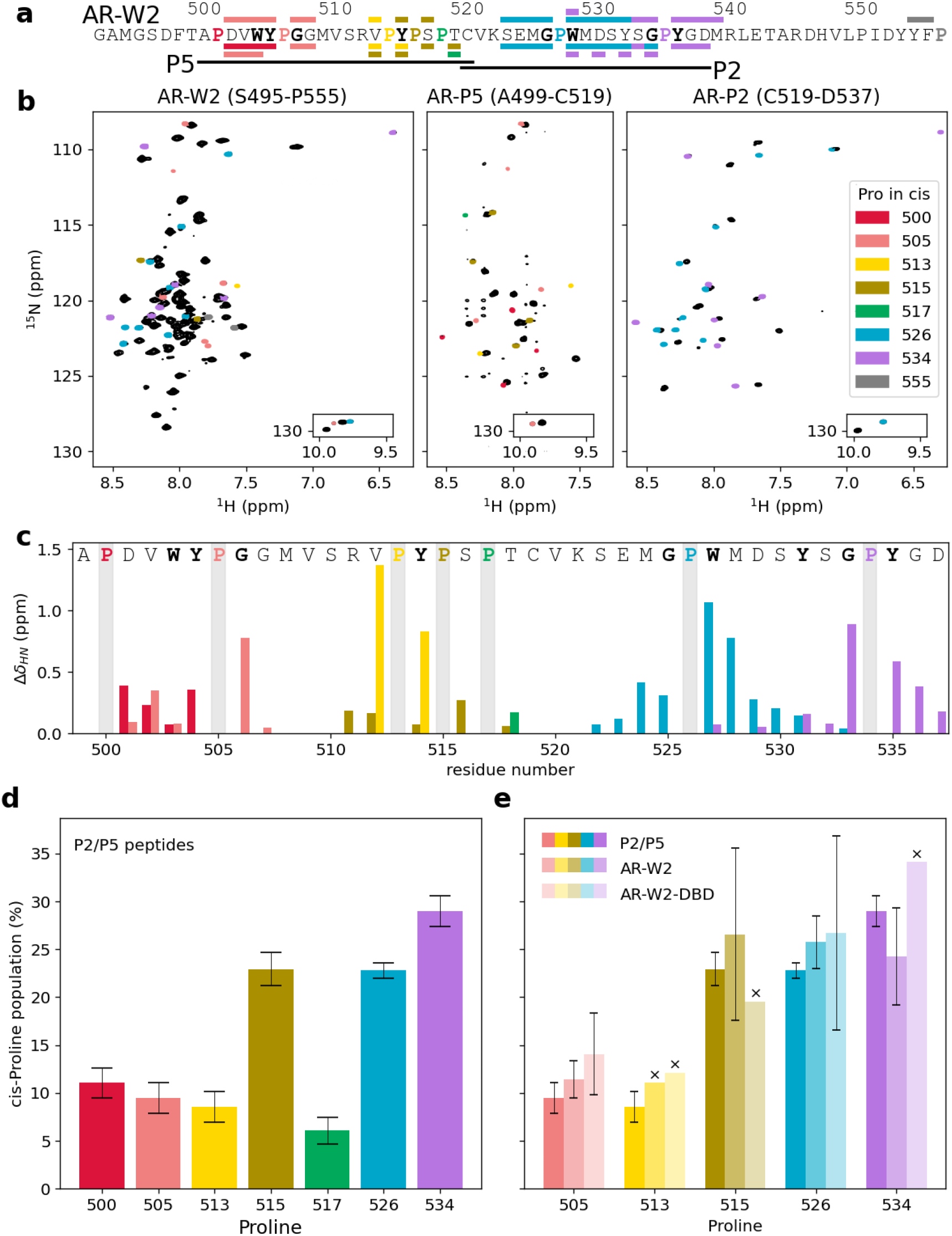
The AR-W2 region is a hotspot of high cis proline populations. The global color coding for individual cis proline conformations is displayed on panel b. **a.** AR-W2 sequence. Regions corresponding to the P5 and P2 peptides are shown as black lines. Prolines with assigned cis populations are shown in colors and coloured bars above and below the sequence show residues whose minor conformation could unambiguously be assigned to the cis conformation of a specific proline in AR-W2 and P5/P2 peptides respectively. **b**. 2D ^1^H-^15^N HSQC spectra of AR-W2 (left), P5 (centre) and P2 (right) correlations assigned to a cis proline population are shown in colour. P5 and P2 spectra were recorded at natural abundance. **c**. Composite H^*N*^ and ^15^N chemical-shift perturbations (Δ*δ*_*HN*_) between cis and trans proline populations in P2/P5 peptides. **d**. cis proline populations for each of the prolines within P5 and P2 peptides. **e**. Comparison of cis proline populations in the P5/P2 peptides (darkest shade), AR-W2 (middle shade) and AR-W2DBD (lighter shade). Crosses indicate prolines for which a single set of peak was used for population estimation because of spectral overlap in longer AR fragments.

The relative populations of trans and cis proline isomers were estimated by comparing the volumes of correlation peaks corresponding to the same nuclei in distinct conformations associated with different proline isomers, using the ^1^H-^15^N and ^1^H-^13^C HSQC spectra for P5/P2 peptides (Figures S5 and S6) and the ^1^H-^15^N HSQC spectra for AR-W2 and AR-W2-DBD. Final proline isomer populations were obtained by averaging the populations derived from all neighbouring nuclei exhibiting more than one set of correlation peaks attributable to proline isomerization (Figure 2d, Supplementary Table 1). This quantitative analysis revealed that cis proline populations critically depend on local sequence context. Prolines could be categorized into two groups based on their isomers populations: low to intermediate cis populations (5–10%) for P500, P505, P513 and P517 and high cis populations (20–30%) for P515, P526 and P534 (Figure 2d). Most minor states assignments could be transferred to longer AR-W2 and AR-W2DBD spectra, thus validating the use of peptide models to study proline isomerisation. Consistent cis proline populations were measured in AR-W2, except for P500/P517, undetectable, possibly due to spectral overlap or reduced cis population in AR-W2 due to edge effects as P500 and P517 were only 1 and 2 residues away from termini in peptides. Besides, in AR-W2, two peaks were assigned to minor states of residues Y553 and F554 for the C-terminal P555 of AR-W2 (not included in the P5/P2 peptides) in cis conformation (27%). Comparison with the AR-W2DBD fragment demonstrated that the presence of the DBD does not significantly alter cis proline populations for P505, P513, P515, P526 and P534 (Figure 2e). This observation suggests that local sequence context, rather than long range interactions notably with the DBD, governs proline isomerization in AR-W2, at least in the absence of DNA or other binding partners. Therefore the P2 and P5 peptides provide good models to characterise proline isomerisation in AR-W2. We next investigated how specific sequence motifs dictate these conformational preferences.

### 2.3 Gly-Pro-Aro motifs favor cis-proline conformations in AR-NTD

Out of the 9 prolines of AR-W2, 6 prolines have adjacent aromatic residues (i+/-1 position): P505, P513, P515, P526, P534 and P555 and 3 prolines have no aromatic neighbors: P500, P517, P549. All 6 aromatic-neighboring prolines showed cis populations higher than 10% in AR-W2, while the 3 non-aromatic prolines displayed sparsely populated cis states ( ≤10% in peptides for P500 and P517 and undetectable cis population in AR-W2 for P500, P517 and P549). This is in line with previous observations of high cis proline populations in presence of aromatic neighbours (Sebák et al., 2023) and supports a model where aromatic-proline CH-*π* interactions stabilize cis conformations via *π*-stacking onto the pyrrolidine ring (Wu and Raleigh, 1998; Ganguly et al., 2013; Zondlo, 2013; Ganguly et al., 2025).

Among aromatic-neighbouring prolines, three exhibited a particularly high cis proline population, over 20%: P515, P526 and P534. P515 is the central proline of the proline-rich VPYPSPT motif where Y514 is sandwiched between P513 and P515. On the other hand, prolines P526 and P534 are embedded in similar Gly-Pro-Aro motifs (GPW and GPY respectively) and both exhibited remarkably high cis populations and extended range of chemical shift perturbations in adjacent residues, up to 12 residues for P526. Those high cis proline populations are stable for temperatures ranging from 5 to 35 degrees celsius (Figure S7).

Remarkably, the combination of P526 and P534 cis/trans isomerization generated four distinct conformational states, each detectable via unique chemical shift signatures for residues D529, S530 and Y531 (Figure 3). Quantification of these states revealed a cis-cis conformer (8% population), consistent with the product of individual cis probabilities (P526: 23% cis; P534: 29% cis; expected cis-cis: 6.6%). Besides chemical shift perturbations observed for the cis-cis conformer compared to the trans-trans are the sum of the perturbations for cis-trans and trans-cis conformers. These observations suggest that prolines P526 and P534 isomerize largely independently, with minimal cooperative interactions.

**Figure 3.**
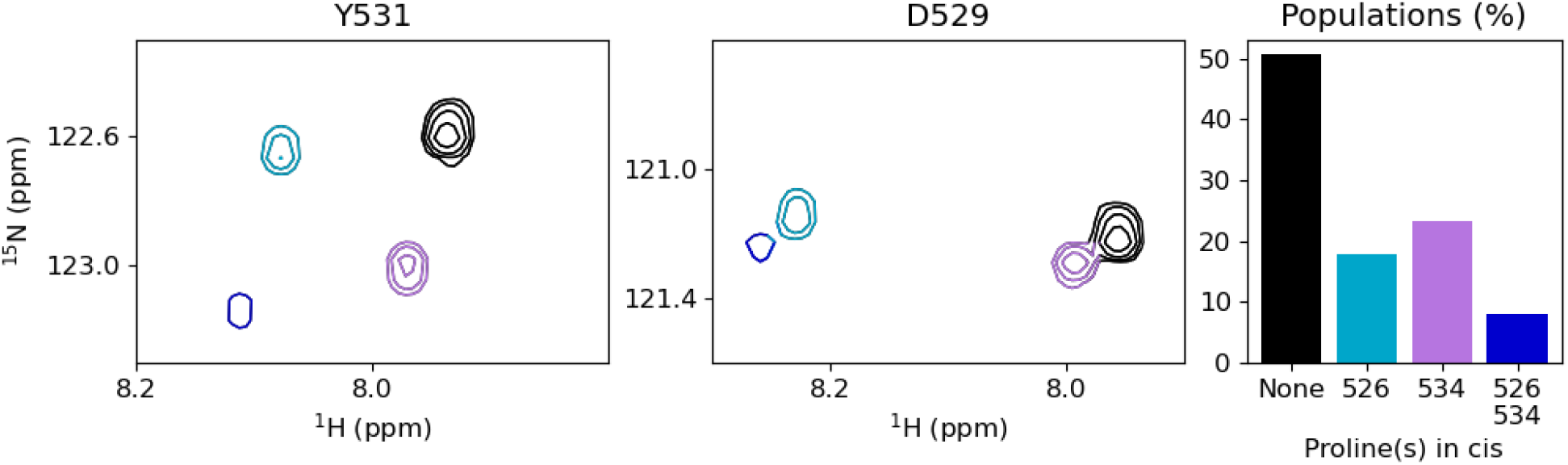
Four combinations of P526 and P534 trans and cis conformations observed within the P2 peptide. Zoomed in regions of the ^1^H-^15^N HSQC spectrum of P2 showing for Y531 (left), D529 (middle) the combinaisons of proline isomers: tP526/tP534 (black), cP526/tP534 (cyan), tP526/cP534 (purple) and cP526/cP534 (navy). Populations of isomers were quantified using the peak volumes of these 2 residues (right).

Interestingly, the high cis populations of P526 and P534 in the context of a Gly-Pro-Aro motif contrast with the lower cis population observed for P505 embedded in a reversed Aro-Pro-Gly motif (between 9.5 and 14.1% in peptide P5, W2 and W2DBD fragments). This motif is part of a longer Aro-Aro-Pro-Gly-Gly sequence (WYPGG) so in order to discard the potential effects of more distant sequence context, we synthesised the P2inv peptide where the GPW motif of P526 is converted into WGP. Thus, we took advantage of the nearly palindromic sequence: _523_S-MGPWM-S_530_ surrounding the GPW of P526 to investigate the effect of the relative position of the glycine and aromatic residue surrounding the proline by comparing cis proline populations in P2 and P2inv. The population of cis-P526 dropped from 22.8 ± 0.8 to 17.2 ± 3.0% suggesting that the Gly-Pro-Aro motif better stabilizes cis proline conformations than Aro-Pro-Gly independently of the more distant sequence context (Figure S8).

### 2.4 The position of proline-adjacent aromatic residues affects *π* **stacking and rotamers differently for cis and trans proline conformers**

To understand the stabilisation of cis proline conformations in these Gly-Pro-Aro motifs, we analysed distance restraints provided by NOESY experiments. NOE peaks were observed between the aromatic protons of W527 and Y535 and protons of the preceding proline in i-1, as well as with the H_*α*_ of the Glycine in i-2, regardless of whether the proline adopted a cis or trans conformation. These observations are consistent with CH-*π* interactions between the aromatic cycle and the pyrrolidine ring of the proline or with the H_*α*_ of the preceding glycine.

Chemical shift analysis of ^1^H-^15^N and ^1^H-^13^C HSQC spectra further revealed significant differences between the cis proline and trans proline conformations in the Gly-Pro-Aro motifs. Whithin these motifs, the H_*α*_ of G525 and G533 (i-1 of P526 and P534 respectively) exhibited upfield shifts of 0.6–1.3 ppm in the cis conformation compared to the trans conformers (Figure S9). Additionally, the H_*N*_ chemical shift of G533 was shifted upfield by 0.8 ppm when P534 adopted a cis conformation compared to the trans conformer (Figure S9). These observations are consistent with pronounced ring current effects arising from the stacking of neighboring aromatic side chains (W527 and Y535) onto the glycines preceding the proline with a Gly-Pro cis peptide bond.

Conversely, in the Aro-Pro-Gly motifs that encompass P505 in peptide P2 (YPG) and P526 in peptide P2inv (WPG), the chemical shift of prolines H_*α*_, H_*β*_ and to a lesser extent H_*γ*_ and H_*δ*_were shifted upfield in the cis conformation compared to in the trans conformation (Figure S10). This suggests that for this reverse motif, the aromatic side chain stacks preferentially on the proline pyrrolidine ring when the Aro-Pro peptide bond is in cis conformation.

Besides, in Gly-Pro-Aro motifs, the geminal *β* -protons of aromatic residues exhibited distinct behaviours between cis and trans proline conformers. For prolines P526 and P534 in the trans conformation, the H_*β*_ chemical shifts of W527 and Y535 were either degenerated or displayed only small chemical shift differences between geminal protons. By contrast, in the cis conformation, the chemical shift difference between geminal H_*β*_ increased, consistently with reduced dynamics of the aromatic side chain. Conversely, in Aro-Pro-Gly motifs (e.g., WPG in P2inv P526 and YPG in P5 P505), the geminal H_*β*_ of aromatic side chains exhibited large chemical shift differences for trans proline populations, while these differences were attenuated or null for cis proline populations (Figure 4, Figure S11).

**Figure 4.**
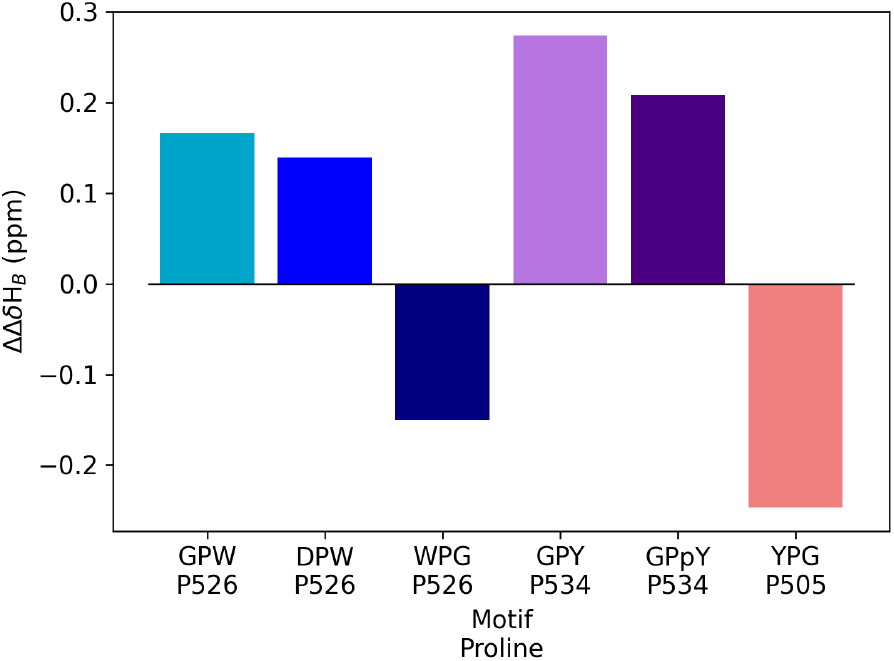
Difference in chemical shift between the geminal H_*β*_ of proline-adjacent aromatic residues in cis vs. trans proline conformations. pY, phosphorylated tyrosine.

Thus, while aromatics adjacent to prolines interact with the proline ring in both cis and trans conformations, the dynamics of the aromatic side chain are reduced when the aromatic residue is at position i+1 of a cis proline or at position i-1 of a trans proline. This pattern may reflect more stable CH-*π* interactions in these conformations, potentially explaining the higher populations of Gly-Pro-Aro motifs in AR-W2.

To further probe the conformations of proline-adjacent aromatic side chains, we measured the intensity ratio of H_*N*_/H_*β*_ NOE peaks for residues with two distinct *β* -protons. This ratio enables assessment of whether the time-averaged H_*N*_/H_*β*_ distance is similar for the two geminal *β* -protons. In a disordered protein, where side chain dynamics are expected to occur on timescales much faster than the NOESY mixing time (200 ms), this ratio can quantify imbalances in rotamer populations. All mesured ratios were above 0.65 (Table 1), indicating relative balance of rotamer populations.

**Table 1.** Ratio of H_*N*_/H_*β*_ NOE peak intensities for the two H_*β*_ of the aromatic side chain. No value is reported when the two H_*β*_ chemical shifts are degenerated. pY, phosphorylated tyrosine.

| Motif | Proline | Aromatic | $(H_N/H_\beta)$ ratio | |
| --- | --- | --- | --- | --- |
|  |  |  | transP | cisP |
| GPY | P534 | Y535 | 0.84 | 0.66 |
| GPpY | P534 | pY535 | 0.68 | 0.66 |
| GPW | P526 | W527 | - | 0.96 |
| WPG | P526 | W525 | 0.76 | - |
| YPG | P505 | Y504 | 0.65 | - |

We observed that, for Y535, the rotamer population was more biased in the cis P534 conformation (0.66) than in the trans P534 conformation (0.84). This bias may arise from favorable CH-*π* interactions between the aromatic ring and the backbone of G533 in the cis P534 conformation. However, the ratio for W527 in the GPW motif (cis P526 conformation, 0.96) was less biased, suggesting that rotamer populations alone cannot fully account for the preferential stabilization of the cis Gly-Pro-Aro motif.

### 2.5 PTM or mutations modulate cis proline populations in AR-W2

Because both Gly-Pro-Aro motifs of P526 and P534 in AR are known to undergo post-translational modifications or to harbour pathological mutations, we next examined how such modifications influence proline cis/trans conformation in these motifs.

Glycine 525 (part of the GPW motif) was found to be mutated to an aspartate in prostate cancer patients (Hyytinen et al., 2002). This mutation was shown to reduce AR sumoylation at K521 and was associated to increase transcriptional activity at genes containing multiple androgen response elements (Mukherjee et al., 2012). Introduction of the G525D mutation in the P2 peptide induced a strong decrease in cis population of P526 (Figure 5a, Figure S12). Chemical shift perturbations induced by the G525D mutation are more pronounced when the neighbouring proline 526 is in cis conformation of this peptide bond compared to the two other states when P526 is in trans. Mobility of aromatic side chain of W527 present in Pro i+1 positions is not affected by the mutation, as evidenced by similar geminal H_*β*_ chemical shift differences for W527 in the G525D mutant (Figure 4, Figure S11). This suggests that the decrease in P526 cis population is mainly due to steric hindrance of the larger aspartate in i-1 position compared to the wild type glycine rather than modulation of interactions of W527 aromatic side chain.

**Figure 5.**
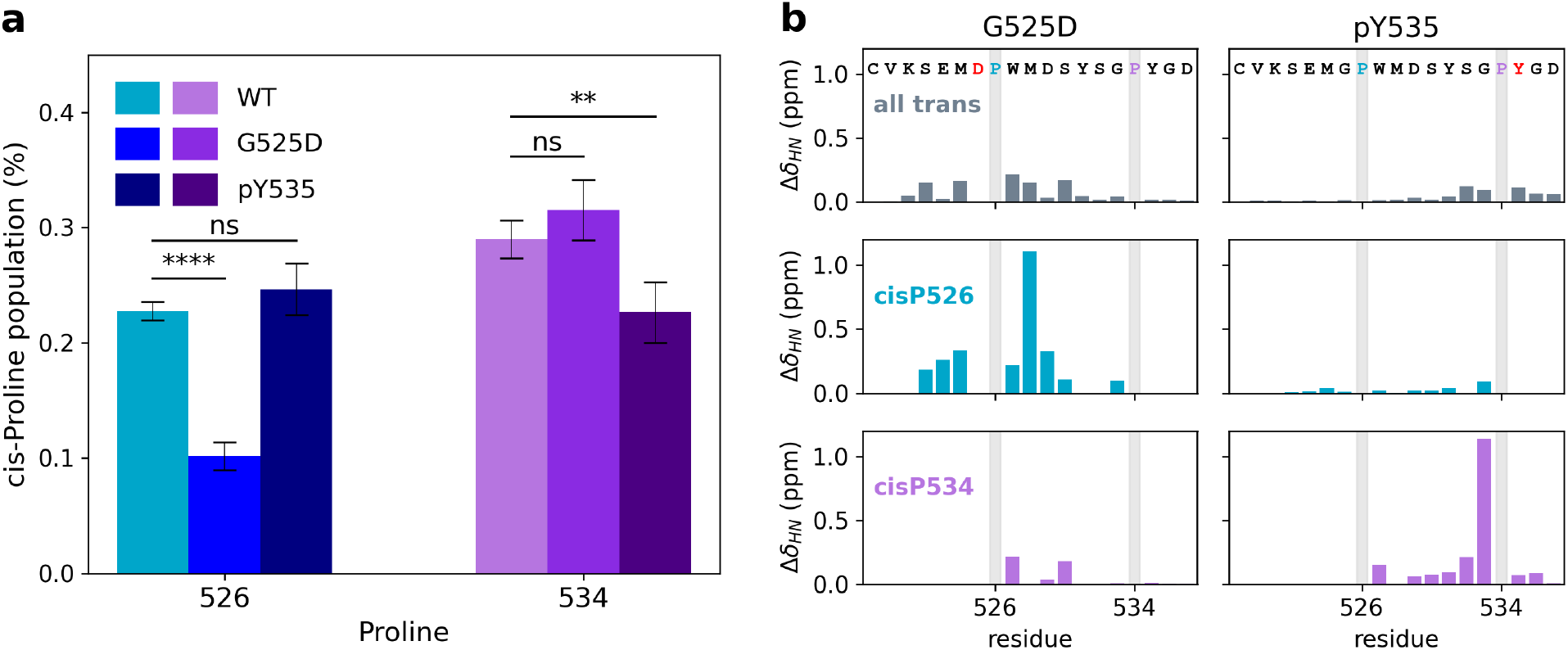
Modifications in the Gly-Pro-Aro motifs of AR-W2 modulate cis proline populations. **a.** cis proline populations for each of the prolines within WT (left), Y535 phosphorylated (middle) and G525D (right) P2 peptides. **b**. Weighted chemical shift perturbations between WT and modified P2 peptides for the fully trans proline cisP526 and cisP534 conformers.

Tyrosine 535 (part of the GPY motif) is a phosphorylation site by Src or Etk kinases (Guo et al., 2006; Dai et al., 2010). Y535 phosphorylation was shown to increase AR stability, nuclear translocation, transcriptional activity and to promote cell growth under androgen depleted conditions (Guo et al., 2006). To elucidate the structural consequences of this modification, we investigated the effect of tyrosine 535 phosphorylation in the context of the P2 peptide. Phosphorylation of the P2 peptide resulted in a selective reduction in the cis population of P534 from 29.0 ± 1.6% to 22.6 ± 2.6%, while the cis population of P526 remained similar, from 22.8 ± 0.8% to 25 ± 2.2% (Figure 5a, Figure S13). Decrease in cis P534 is accompanied by chemical shift perturbations in the phosphorylated vs WT P2 ^1^H-^15^N HSQC spectra (Figure 5b). The most significant change was observed for the amide proton of G533, which shifts by approximately 1ppm upon phosphorylation of Y535 for the cis isomer of the Gly-Pro peptide bond of P534. This dramatic effect suggests that the ring current effects of the Y535 aromatic side chain on G533 are attenuated upon Y535 phosphorylation (Figure S9). In addition, phosphorylation reduced the chemical shift difference between the geminal H_*β*_ protons of Y535 exclusively in the cisP534 conformer, while the corresponding shifts for Y535 in the transP534 conformer and W527 in the cisP526 conformer remained unaffected (Figure 4, Figure S11). Overall, the cisP534 conformer exhibited greater perturbations than the transP534 conformer, highlighting a more pronounced conformational impact on the cis state. Finally, although the H_*N*_/H_*β*_ NOE ratios for phosphorylated Y535 in the cisP534 conformer were unchanged relative to the unphosphorylated peptide, this ratio decreased from 0.84 to 0.68 for the transP534 conformer (comparing Y535 and pY535, respectively), indicating a significant alteration in rotamer populations for the transP534 state upon phosphorylation (Table 1).

Collectively, these observations suggest that Y535 phosphorylation increases the flexibility of its side chain, a result of the potential weakening of its interactions with the neighbouring proline in cis conformation. Thus tyrosine phosphorylation in the Gly-Pro-Try motif reduces the population of the cis proline conformer.

## 3. Discussion

We characterised the region spanning the last 65 residues of AR-NTD (residues S495–P555) named AR-W2 for its 2 conserved tryptophans (W503 and W527). This highly conserved region exhibits an unusual composition for an intrinsically disordered region, with a high abundance of aromatic and nonpolar residues and a remarkable enrichment in prolines with 9 prolines distributed across the sequence (15% of total residues). NMR characterisation of several fragments derived from this region revealed that this intrinsically disordered region displays extensive conformational heterogeneity linked to Xaa-Pro peptide bond cis/trans isomerisation. Three sites, P515, P526 and P534, were identified as cis proline hotspots, with cis proline populations superior to 20%, significantly higher than the ≤10% cis proline baseline observed in most IDRs (Alderson et al., 2018; Mateos et al., 2020).

Aromatic residues in i+/-1 position relative to prolines were previously observed to stabilise cis proline conformations (Mateos et al., 2020; Sebák et al., 2023), possibly through CH-*π* interactions between the aromatic and proline pyrrolidine rings (Wu and Raleigh, 1998; Zondlo, 2013; Ganguly et al., 2013; Ganguly et al., 2025). Within AR-W2, the Gly-Pro-Aro motifs including P526 and P534 exhibited higher cis proline populations than the Aro-Pro-Gly motif centred on P505. Our data suggest that the aromatic side chain conformations are more restricted for cis proline conformers when the aromatic is in the i+1 position (Figure 6), whereas they are more restricted in trans conformers when the aromatic is in the i-1 position.

**Figure 6.**
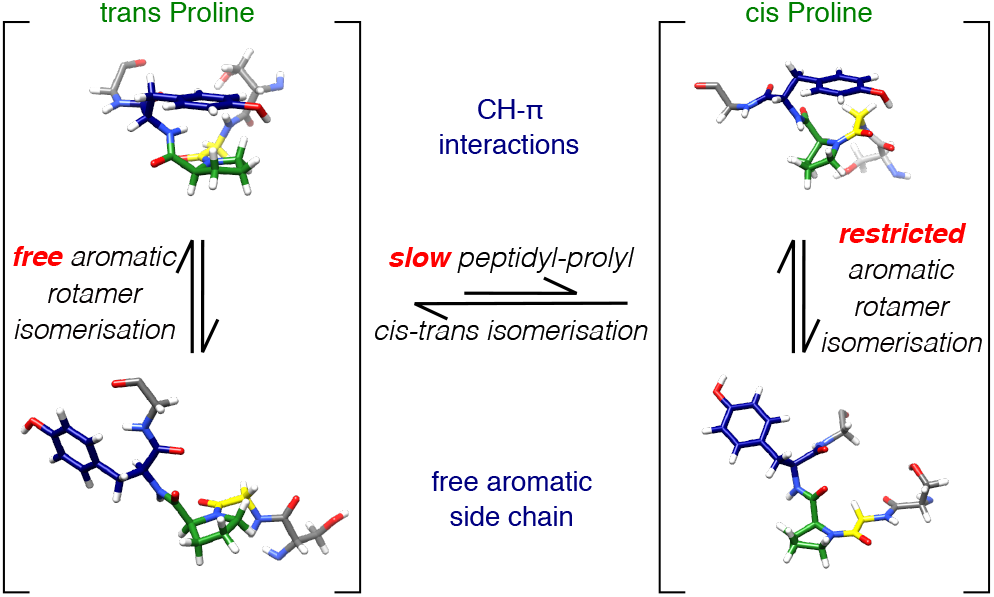
Aromatic proline interaction in Gly-Pro-Aro peptides. The Gly-Pro peptide bond slowly isomerises between cis and trans conformations. The aromatic side chain of the residue following the proline interacts with the proline for both isomers and with glycine in i-1 for the cis proline conformer through dynamic CH-*π* interactions. Example conformations are drawn for a SGPYG peptides where the glycine, proline and tyrosine residues of the Gly-Pro-Aro motif are shown in yellow, green and blue respectively.

Notably, two modifications of the Gly-Pro-Aro motifs are found to affect AR gene activation function, namely the pathological G525D mutation found in prostate cancer and the physiological phosphorylation of Y535. We found that both modifications alter the conformational heterogeneity due to proline cis/trans isomerisation by reducing cis proline populations. Tyrosine phosphorylation decreases electron density in the aromatic ring, as the electron-donating phenol group is converted into a phosphate ester. This would lead to weakening of the CH-*π* interactions of the aromatic side chain with the preceding proline and glycine residues, which could explain increase of aromatic side chain dynamics in the cis proline conformer and the shift of the cis/trans proline equilibrium toward the trans proline conformer. This is consistent with a previous study using non canonical amino acids that links high electron density in aromatic side chains adjacent to prolines to higher cis proline population (Thomas et al., 2006). Thus tyrosine phosphorylation may act as a molecular switch, tuning IDRs conformational ensemble through modulation of cis/trans proline isomer’s population, as serine and threonine phosphorylation are known to favour the adjacent trans proline conformer, although through different mechanisms.

Other genetic alterations in AR-W2 further emphasize the functional importance of prolines in this region. Among the 15 prostate cancer- or AIS-related missense mutations identified in AR-W2, five result in proline suppression (P505T, P515S, P517S, P534S, and P549S), while another five directly modify proline-adjacent residues.

In highly dynamic intrinsically disordered regions, prolines play a unique role: the slow isomerisation between their cis and trans conformers effectively partitions the conformational ensemble into distinct, slowly interconverting subpopulations, each with unique structural and possibly functional properties. High cis proline populations coupled with their cis/trans isomerisation dynamics have been identified as critical regulatory elements in diverse biological processes, including NCBD-ACTR interaction (Zosel et al., 2018), 14-3-3 complex formation (Theisen et al., 2025), DREB2A-Med25 interactions in plants stress response (Theisen et al., 2024) but also for PPAR nuclear receptor transcriptional response (Williams et al., 2025) or Tau liquid liquid phase separation (Zhuang et al., 2024).

The functional outcome depends on the relative populations of each isomer, which can be further modulated by post-translational modifications of proline-adjacent residues (Theisen et al., 2025). Moreover, the slow proline cis/trans isomerisation introduces a kinetic component to signal transduction, acting like a molecular timer. Beyond intrinsic isomerization dynamics, the functional consequences of proline conformations in IDPs are further modulated by prolyl isomerases, which accelerate proline isomerisation and thus influence the kinetic partitioning of cis/trans subpopulations (Williams et al., 2025; Dujardin et al., 2015). In nuclear receptors, interactions between disordered NTD prolines and the proline isomerase Pin1 were shown to modulate transcriptional response of PPAR (Williams et al., 2025), increase DNA binding of ER*α* (Rajbhandari et al., 2015) and increase GR recruitment at target genes (Poolman et al., 2013). In the androgen receptor specifically, Pin1 was observed to interact with prolines at positions 84, 259 and 311 (La Montagna et al., 2012; Leung et al., 2021) and its inhibition was shown to enhance the efficacy of ralaniten, a NTD-targeting drug (Leung et al., 2021). While immunophilins, such as FKBP51/52 and Cyp40 proteins, are established regulators of AR activity within the unliganded cytoplasmic AR complex (Periyasamy et al., 2007; Periyasamy et al., 2010; Ni et al., 2010; Maeda et al., 2022) their potential interaction with the AR-NTD and associated functional implications remains uncharacterised. Therefore while prolyl isomerases, and in particular Pin1, are emerging as key modulators of AR-NTD conformation and function, their precise role in shaping the conformational ensemble, co-regulator recruitment, and liquid-liquid phase separation properties of AR remains to be fully characterized.

## Conclusion

In this study, we showed that the conserved C-terminal region of the androgen receptor disordered N-terminal domain displays extensive conformational heterogeneity linked to proline isomerisation. Two Gly-Pro-Aro motifs exhibit strikingly high cis proline population associated with interaction of the aromatic ring with the glycine residue preceding the proline, together with restricted aromatic side chain rotamer isomerisation in the cis proline conformation. Interestingly, cis proline populations at these motifs are reduced by a cancer associated mutation in one case and tyrosine phosphorylation in the other. Future work should elucidate how the dynamic tuning of these populations, together with the acceleration of cis/trans exchange by proline isomerases, remodels AR-NTD conformational ensembles, dynamics, and interaction networks, thereby modulating androgen receptor function in health and disease. More broadly, this work highlights the unique ability of NMR spectroscopy to provide atomic-resolution insight with exquisite sensitivity to motions spanning a broad range of timescales, from fast local side chain dynamics to slow proline cis/trans isomerisation. Such multi-timescale information will be essential to understand how proline-rich sequence motifs encode functional conformational heterogeneity within the androgen receptor N-terminal domain and other intrinsically disordered regions.

## Supporting information

Sequence alignment of the W2 region of AR-NTD for representative species; Sequence alignment and amino acid bias of the C-terminal region of steroid receptors NTD, AR-W2 and AR-W2DBD ^15^N-^1^H and ^13^C-^1^H HSQC spectra; Population of cis-Proline conformations in P5, P2, AR-W2, AR-W2DBD, P2inv, P2-G525D and P2-pY535; Population of cis-Proline conformations in P2 as a function of temperature; Glycine, proline and aromatic H*β* C*β* regions of ^15^N-^1^H and ^13^C-^1^H HSQC spectra. Tables S1-2 and Figures S1-13.

## Acknowledgments

This work was funded by the ANR projects ARCHAP (ANR-20-CE11-0004-01) and ARIA (ANR-13-BSV5-0013-03) and by Alsace contre le cancer. Financial support from the IR INFRANALYTICS FR2054 for access to high magnetic field spectrometer in IBS (Grenoble, France) is gratefully acknowledged. The authors thank Pascal Eberling for peptides synthesis and Marc-André Delsuc for insightful comments on the manuscript.

## TOC image

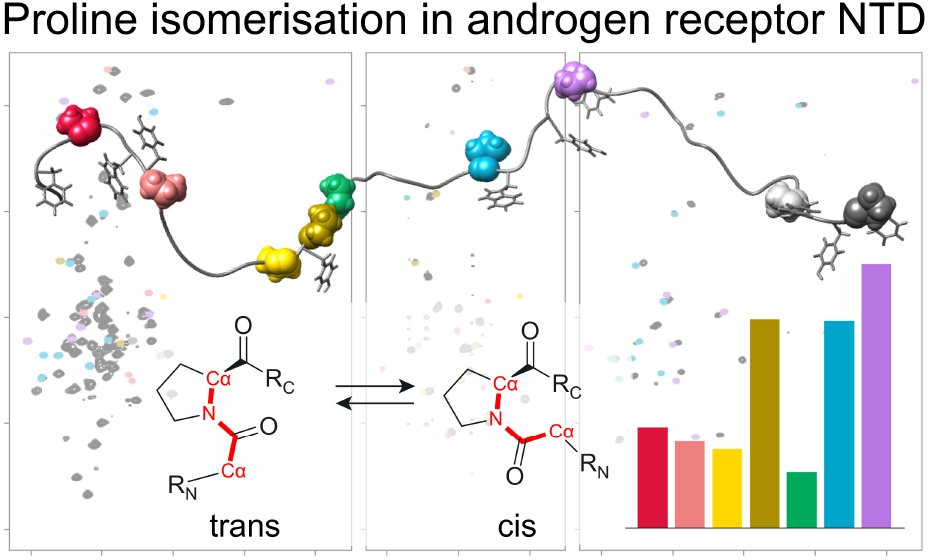

