## Supplementary information for "A hotspot for conformational heterogeneity driven by proline isomerisation in the androgen receptor disordered N-terminal domain"

Mathilde ROTH, Hélène LAUNAY, Eva ERDMANN,
Veronique RECEVEUR-BRECHOT, Jocelyn CERALINE, Bruno KIEFFER and Célia DEVILLE

July 29, 2026

| | P5 | | P2 | | AR-W2 | | AR-W2DBD | |
|---|---|---|---|---|---|---|---|---|
| Proline | cis-Pro (%) | Count | cis-Pro (%) | Count | cis-Pro (%) | Count | cis-Pro (%) | Count |
| 500 | 11.0 ± 1.6 | 16 | | | | 0 | | 0 |
| 505 | 9.5 ± 1.6 | 15 | | | 11.4 ± 1.9 | 7 | 14.1 ± 4.3 | 4 |
| 513 | 8.6 ± 1.6 | 9 | | | 11.1 ± 0.0 | 1 | 12.1 ± 0.0 | 1 |
| 515 | 22.9 ± 1.7 | 21 | | | 26.6 ± 9.0 | 2 | 19.5 ± 0.0 | 1 |
| 517 | 6.1 ± 1.4 | 7 | | | | 0 | | 0 |
| 526 | | | 22.8 ± 0.8 | 37 | 25.8 ± 2.7 | 10 | 26.7 ± 10.2 | 4 |
| 534 | | | 29.0 ± 1.6 | 25 | 24.2 ± 5.1 | 7 | 34.1 ± 0.0 | 1 |
| 549 | | | | | | 0 | | 0 |
| 555 | | | | | 26.6 ± 12.9 | 2 | | 0 |

Table S1: Population of cis-Proline conformations in P5, P2, AR-W2 and AR-W2DBD estimated from relative peak intensities in HN and HC HSQC spectra. The count column indicated the number of correlation peaks used to estimate cis conformation of a given proline.

| | P2inv | | P2-G525D | | P2-pY535 | |
|---|---|---|---|---|---|---|
| Proline | cis-Pro (%) | Count | cis-Pro (%) | Count | cis-Pro (%) | Count |
| 526 | 17.2 ± 3.0 | 9 | 10.2 ± 1.2 | 39 | 24.6 ± 2.2 | 39 |
| 534 | 25.4 ± 5.9 | 7 | 31.5 ± 2.6 | 26 | 22.6 ± 2.6 | 29 |

Table S2: Population of cis-Proline conformations in P2inv, P2-G525D and P2-pY535 estimated from relative peak intensities in HN and HC HSQC spectra. The count column indicated the number of correlation peaks used to estimate cis conformation of a given proline.

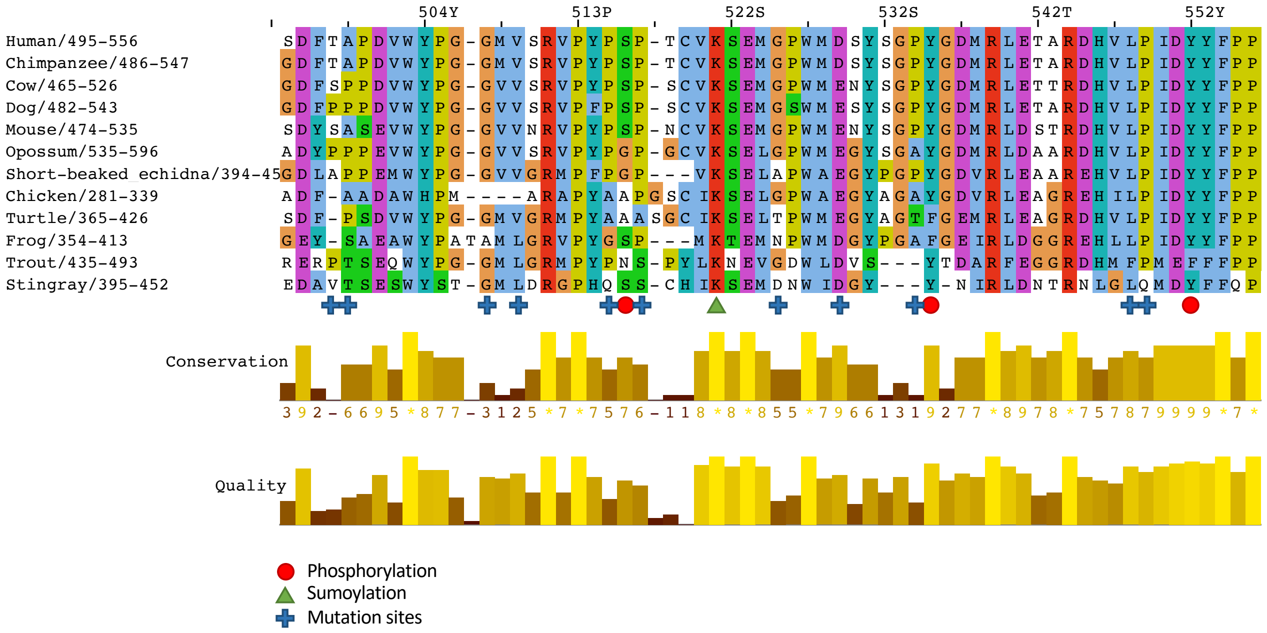


Figure S1: Sequence alignment of the W2 region of AR-NTD for representative species. Phosphorylation, sumoylation and mutation sites are indicated as red disks, green triangle and blue crosses respectively.

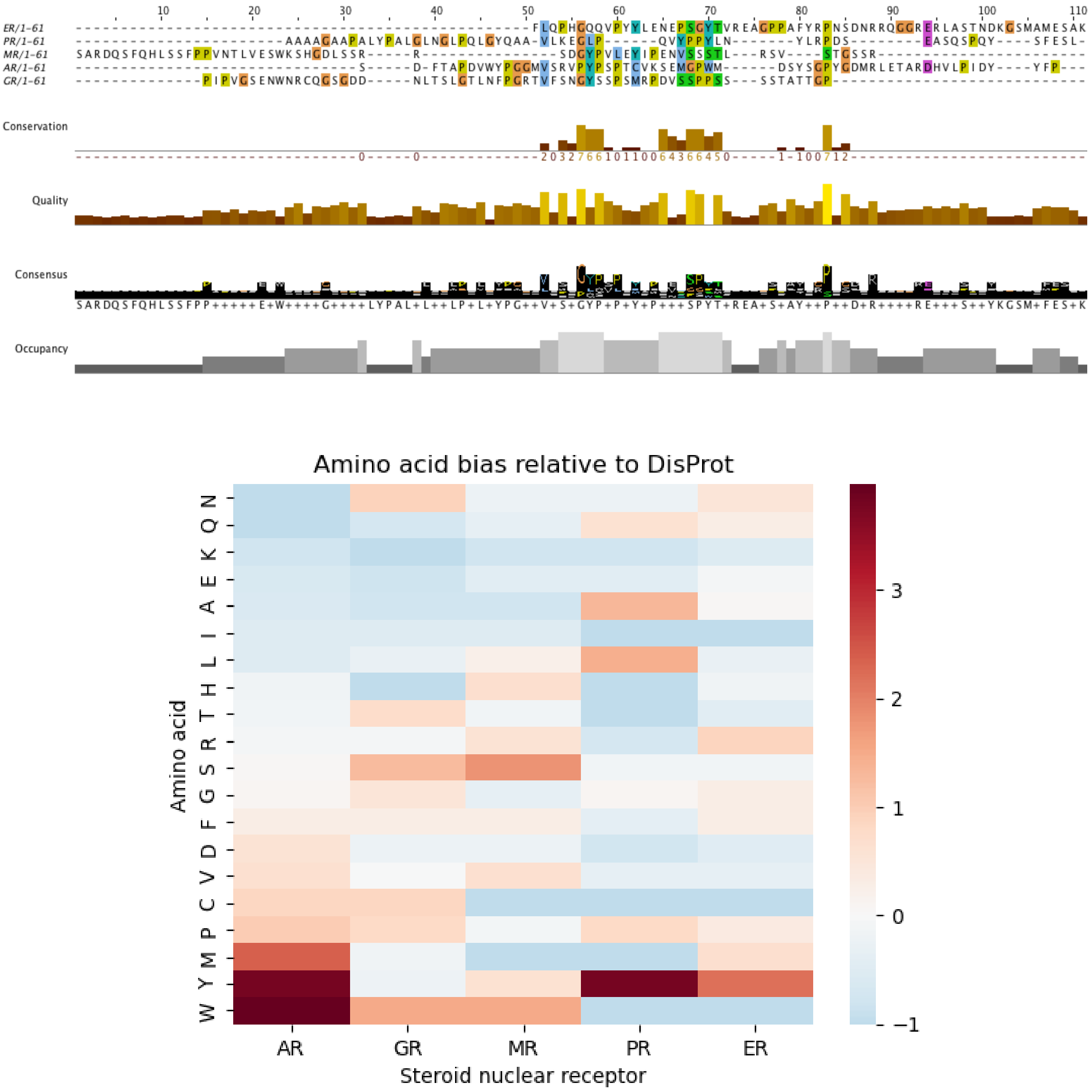


Figure S2: Sequence alignment of the C-terminal region of steroid receptors NTD shows no sequence conservation. Heatmap of amino acid bias in the C-terminal region of steroid receptors NTD shows a very different sequence composition of AR-W2 compared to equivalent regions in other steroid receptors.

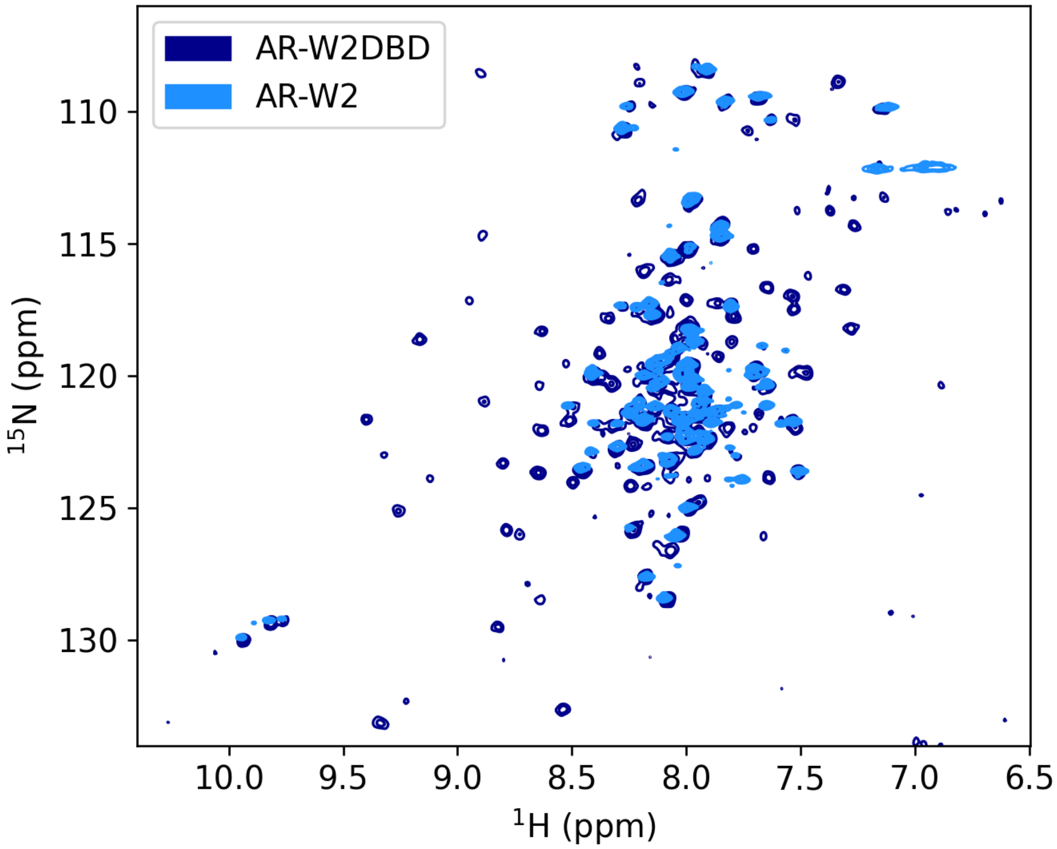


Figure S3: Overlap of AR-W2 and AR-W2DBD $^{15}$N-$^{1}$H HSQC spectra show that AR-W2 remains disordered in presence of the DBD with similar chemical shifts for residues of the W2 region for the 2 fragments.

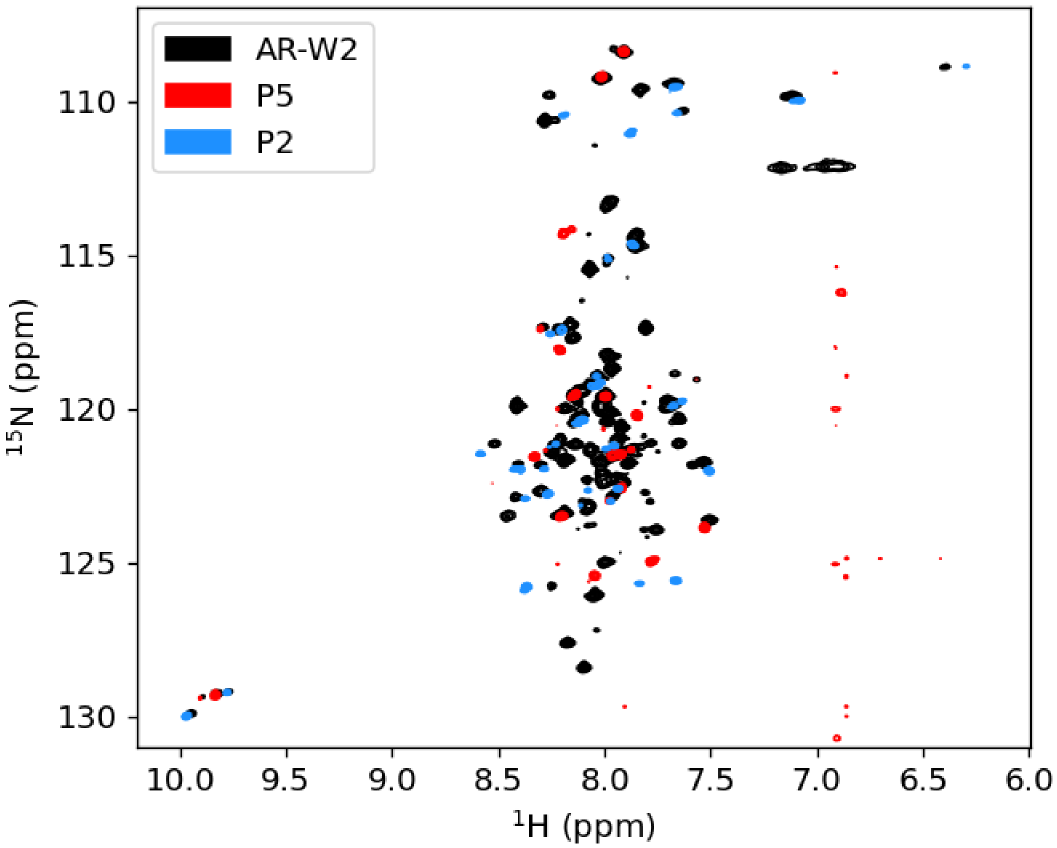


Figure S4: AR-W2 and peptides P5 and P2 $^{15}$N-$^{1}$H HSQC spectra. Resonances of AR-W2 and peptides P5 and P2 overlap except for terminal residues.

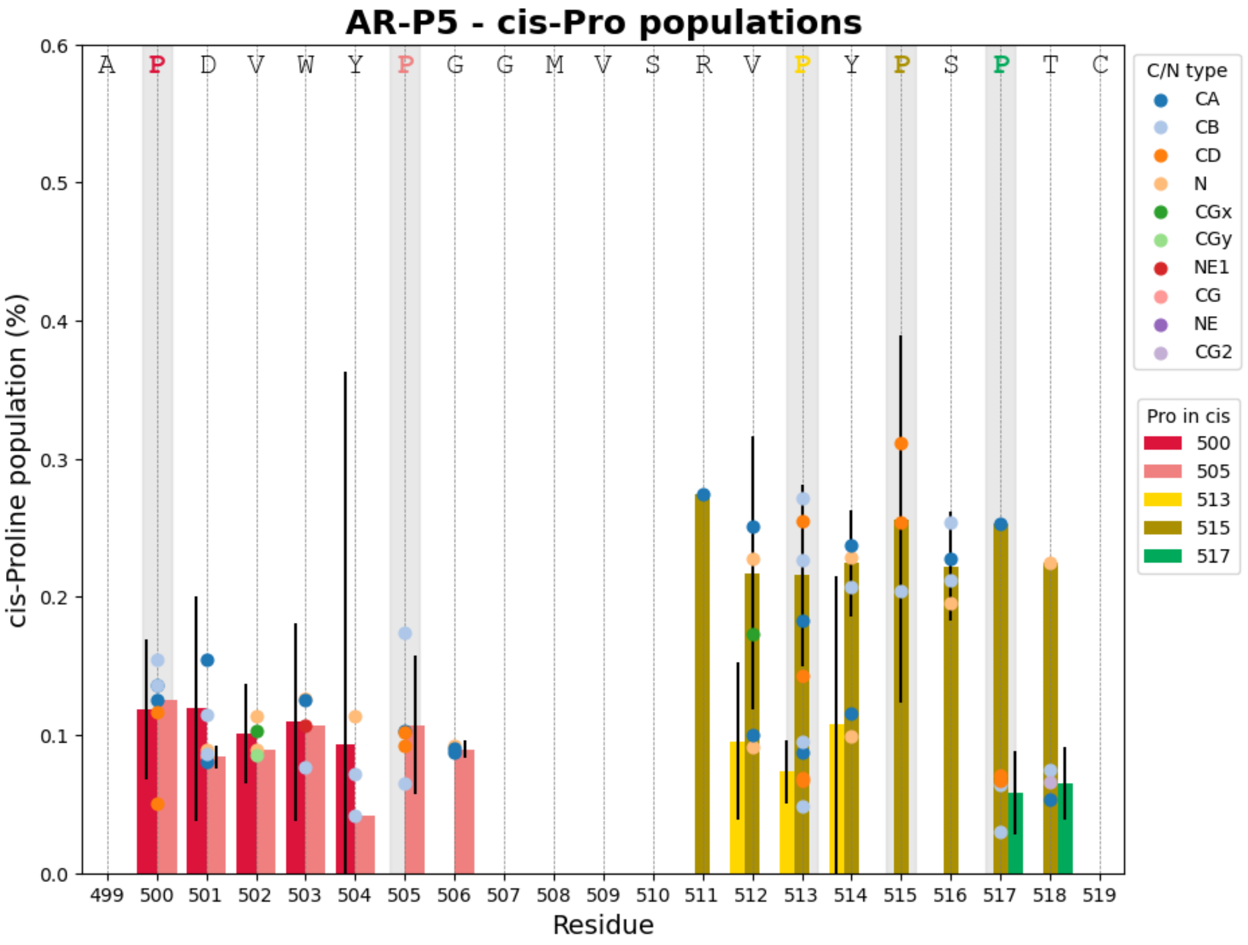


Figure S5: cis-Proline populations for each of the prolines within the P5 peptide. The relative populations of trans and cis proline isomers were estimated by comparing the intensities of correlation peaks corresponding to the same nuclei in distinct conformations associated with different proline isomers, using the $^{15}$N-$^{1}$H and $^{13}$C-$^{1}$H HSQC spectra. Scatter dots show the contribution of individual nuclei. When different correlation peaks could be used for one residue, bars show the average of all nuclei contribution for a given residue and cis Proline conformation. The uncertainty in the population estimates, for one residue and one proline conformation, was calculated as the standard deviation of all nuclei contribution, corrected using the appropriate Student's t coefficient for a two-sided 95% confidence interval. Uncertainty contributions arising from the signal-to-noise ratio of individual peaks were found to be negligible relative to the variability between population estimates obtained from different correlations and were therefore neglected.

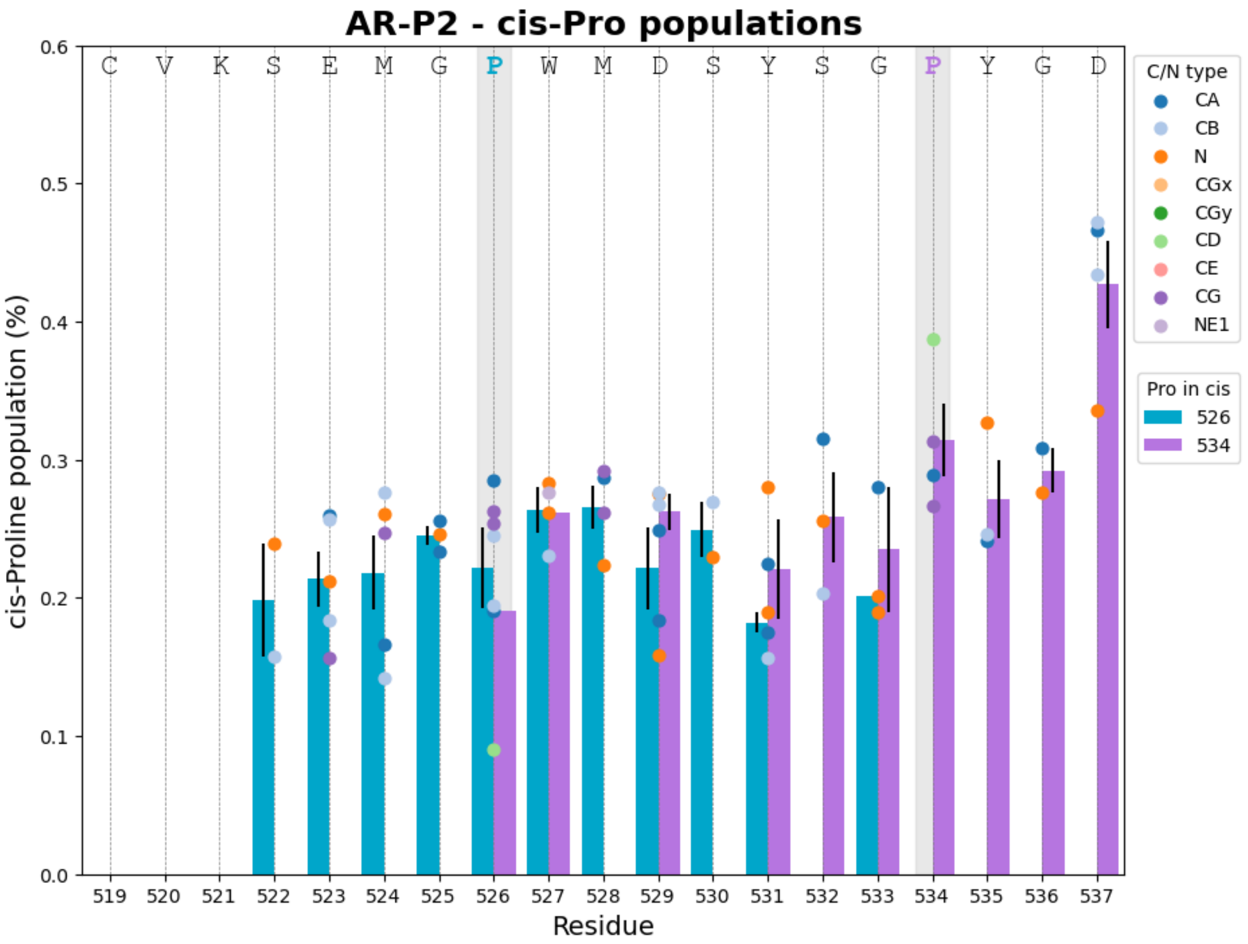


Figure S6: cis-Proline populations for each of the prolines within the P2 peptide. The relative populations of trans and cis proline isomers were estimated by comparing the intensities of correlation peaks corresponding to the same nuclei in distinct conformations associated with different proline isomers, using the $^{15}N$-$^{1}H$ and $^{13}C$-$^{1}H$ HSQC spectra. Scatter dots show the contribution of individual nuclei. When different correlation peaks could be used for one residue, bars show the average of all nuclei contribution for a given residue and cis Proline conformation. The uncertainty in the population estimates, for one residue and one proline conformation, was calculated as the standard deviation of all nuclei contribution, corrected using the appropriate Student's t coefficient for a two-sided 95% confidence interval. Uncertainty contributions arising from the signal-to-noise ratio of individual peaks were found to be negligible relative to the variability between population estimates obtained from different correlations and were therefore neglected.

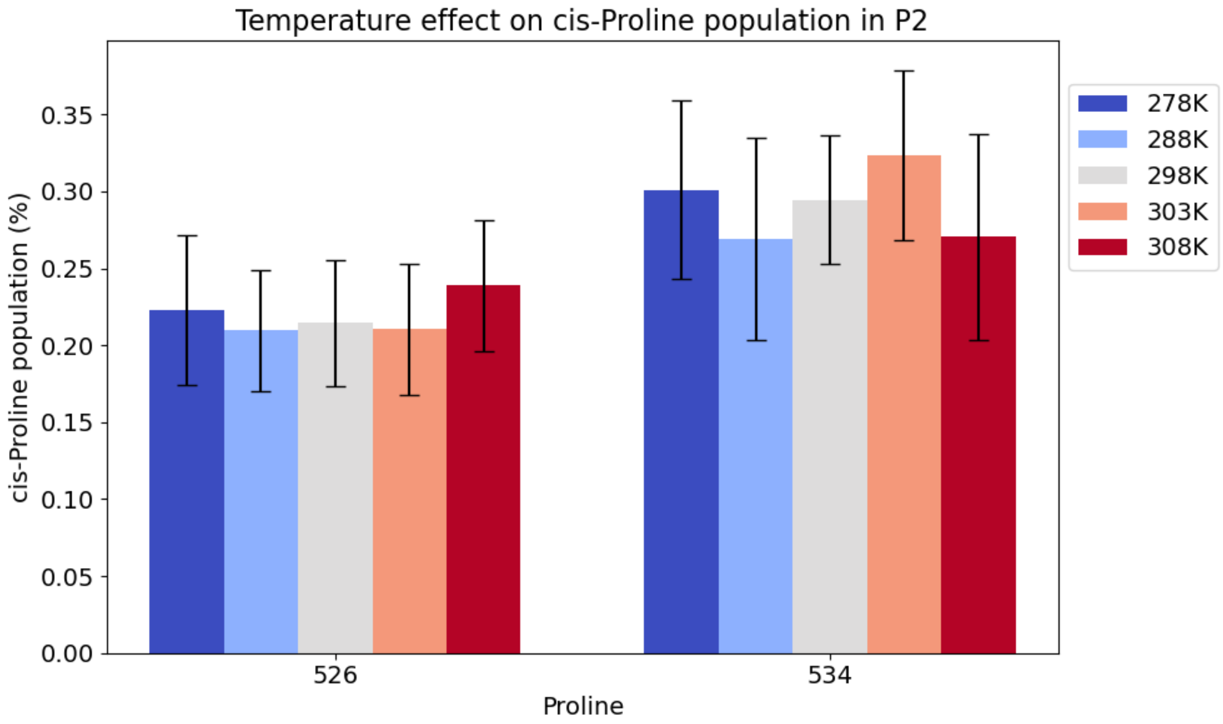


Figure S7: Effect of temperature on proline isomerisation in the P2 peptide. Populations were estimated using peak volumes from the $^{13}$C-$^{1}$H HSQC spectra.

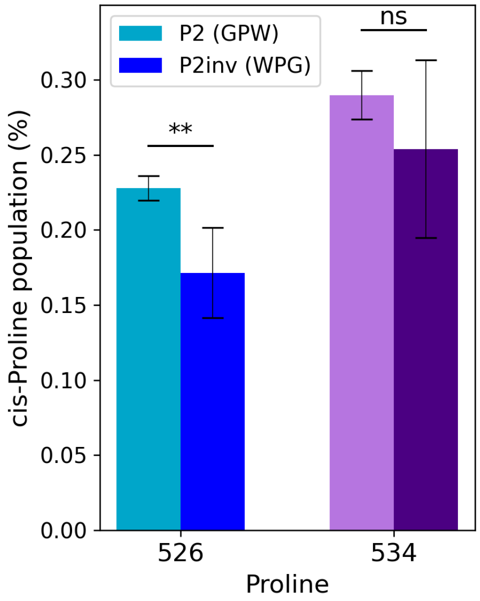


Figure S8: Inversion of the GPW into a WPG motif in the P2 peptide yields a decrease in P526 cis conformer population. Statistical comparison of conformer populations between peptides was performed using an unpaired Student's t test. The corresponding p-values were 0.0025 (**) and 0.2856 (ns) for P526 and P534 cis conformer populations respectively.

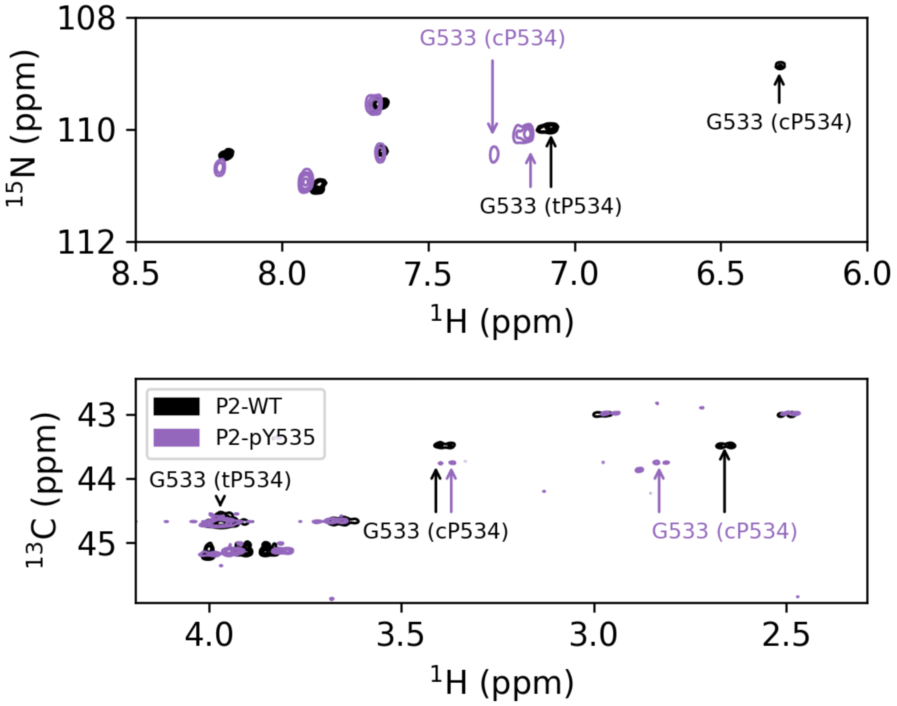


Figure S9: Superimposition of the glycine regions of $^{15}$N-$^{1}$H (top) and $^{13}$C-$^{1}$H (bottom) HSQC spectra for the P2 peptide wild type or including phosphorylation of tyrosine Y535. Y535 phosphorylation leads to loss of shielding of Hn and one of the H$\alpha$ of G533, only for the cisP534 conformer.

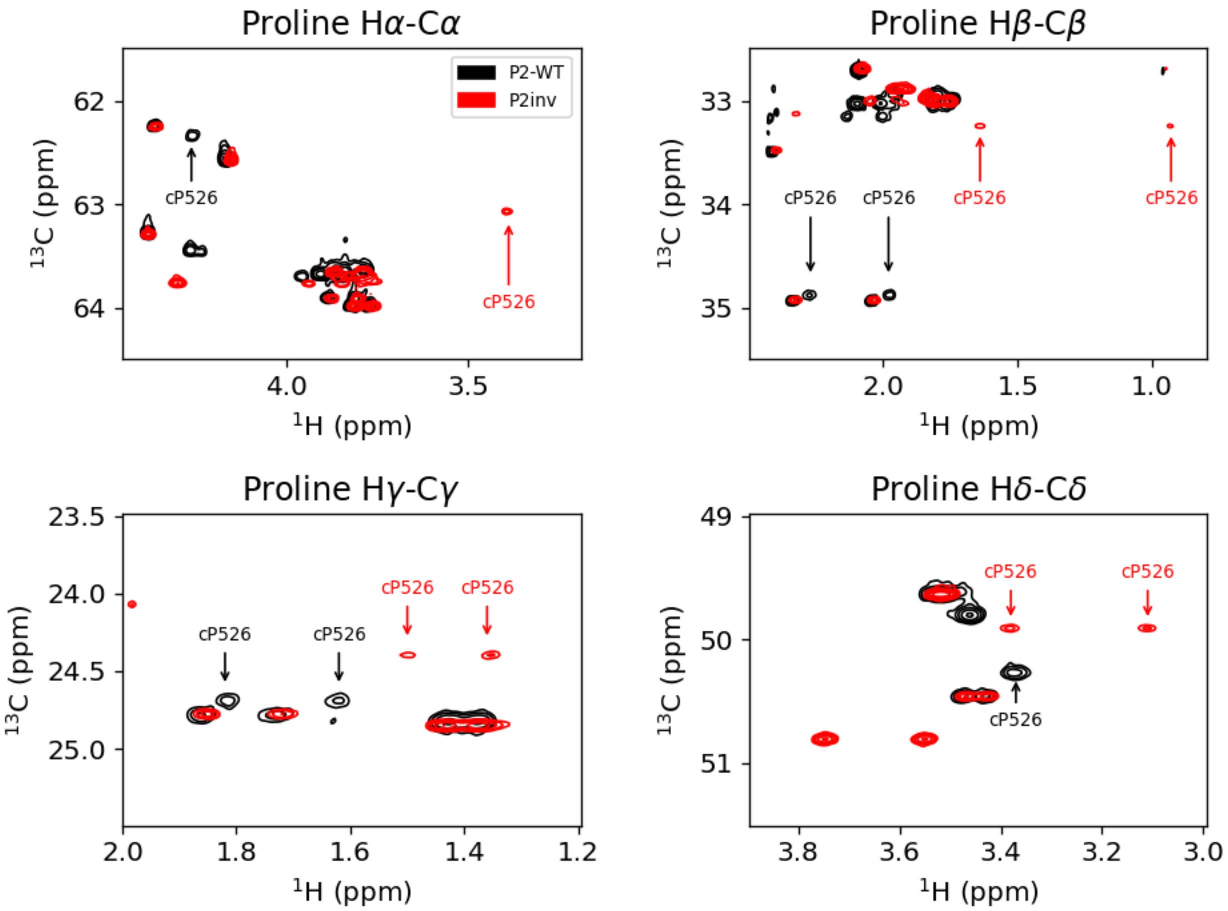


Figure S10: Superimposition of the proline regions of $^{13}$C-$^{1}$H HSQC spectra for the P2 peptide wild type containing a GPW motif at P526 or P2inv including the inverted WGP motif at P526. In P2inv, chemical shift of prolines H$\alpha$, H$\beta$ and to a lesser extent H$\gamma$ and H$\delta$ were shifted upfield in the cis conformation. This suggests that for this reverse motif, the aromatic side chain stacks preferentially on the proline pyrrolidine ring when the Aro-Pro peptide bond is in cis conformation.

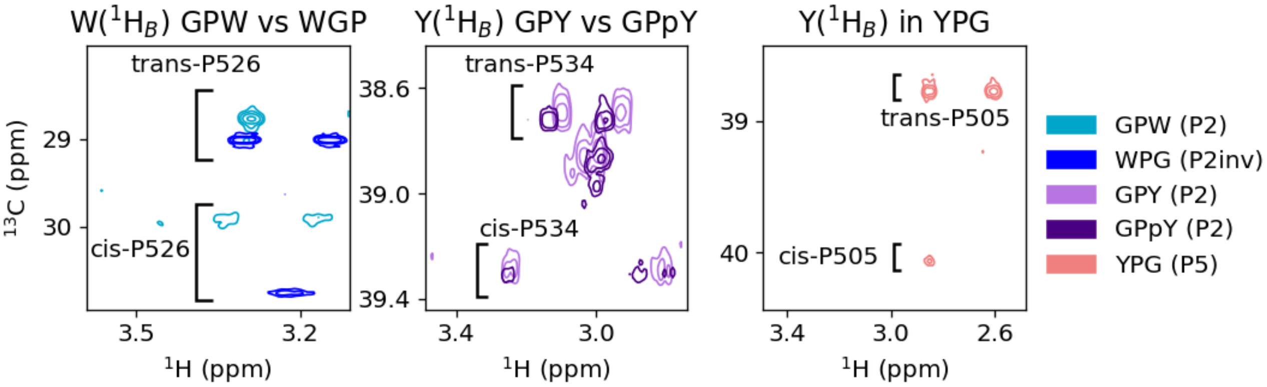


Figure S11: $\beta$-protons of the aromatic residues neighbouring prolines display different behaviour when the neighbouring proline is in cis or trans conformation. H$\beta$C$\beta$ region of the tryptophan (left panel) and tyrosine (middle and right panels) regions show opposite behaviour for Gly-Pro-Aro and Aro-Gly-Pro motifs. Upon phosphorylation of Y535, the chemical shift difference between H$\beta$ geminal protons is reduced.

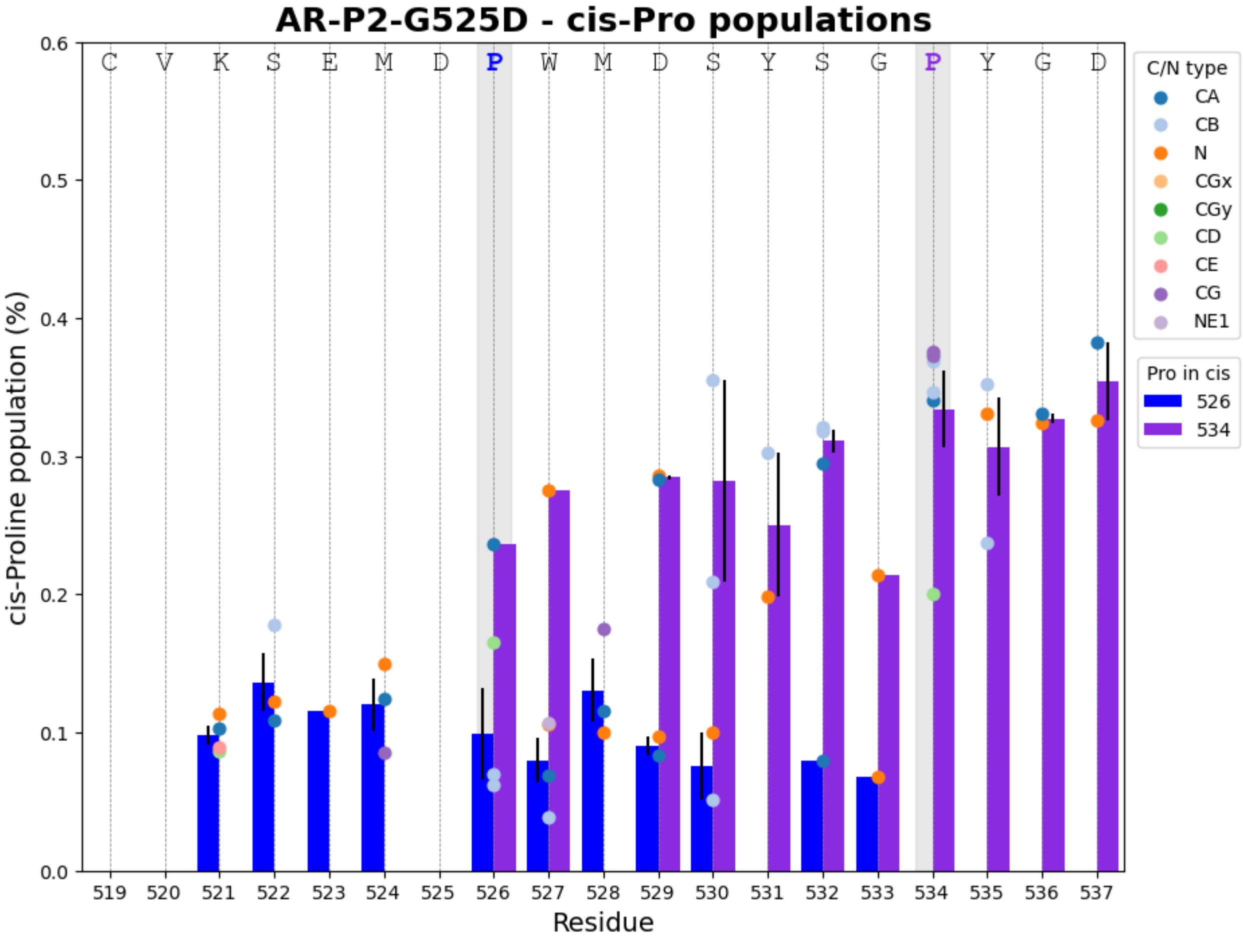


Figure S12: cis-Proline populations for each of the prolines within the P2 peptide arboring the G525D mutation. The relative populations of trans and cis proline isomers were estimated by comparing the intensities of correlation peaks corresponding to the same nuclei in distinct conformations associated with different proline isomers, using the $^{15}$N-$^{1}$H and $^{13}$C-$^{1}$H HSQC spectra. Scatter dots show the contribution of individual nuclei. When different correlation peaks could be used for one residue, bars show the average of all nuclei contribution for a given residue and cis Proline conformation. The uncertainty in the population estimates, for one residue and one proline conformation, was calculated as the standard deviation of all nuclei contribution, corrected using the appropriate Student's t coefficient for a two-sided 95% confidence interval. Uncertainty contributions arising from the signal-to-noise ratio of individual peaks were found to be negligible relative to the variability between population estimates obtained from different correlations and were therefore neglected.

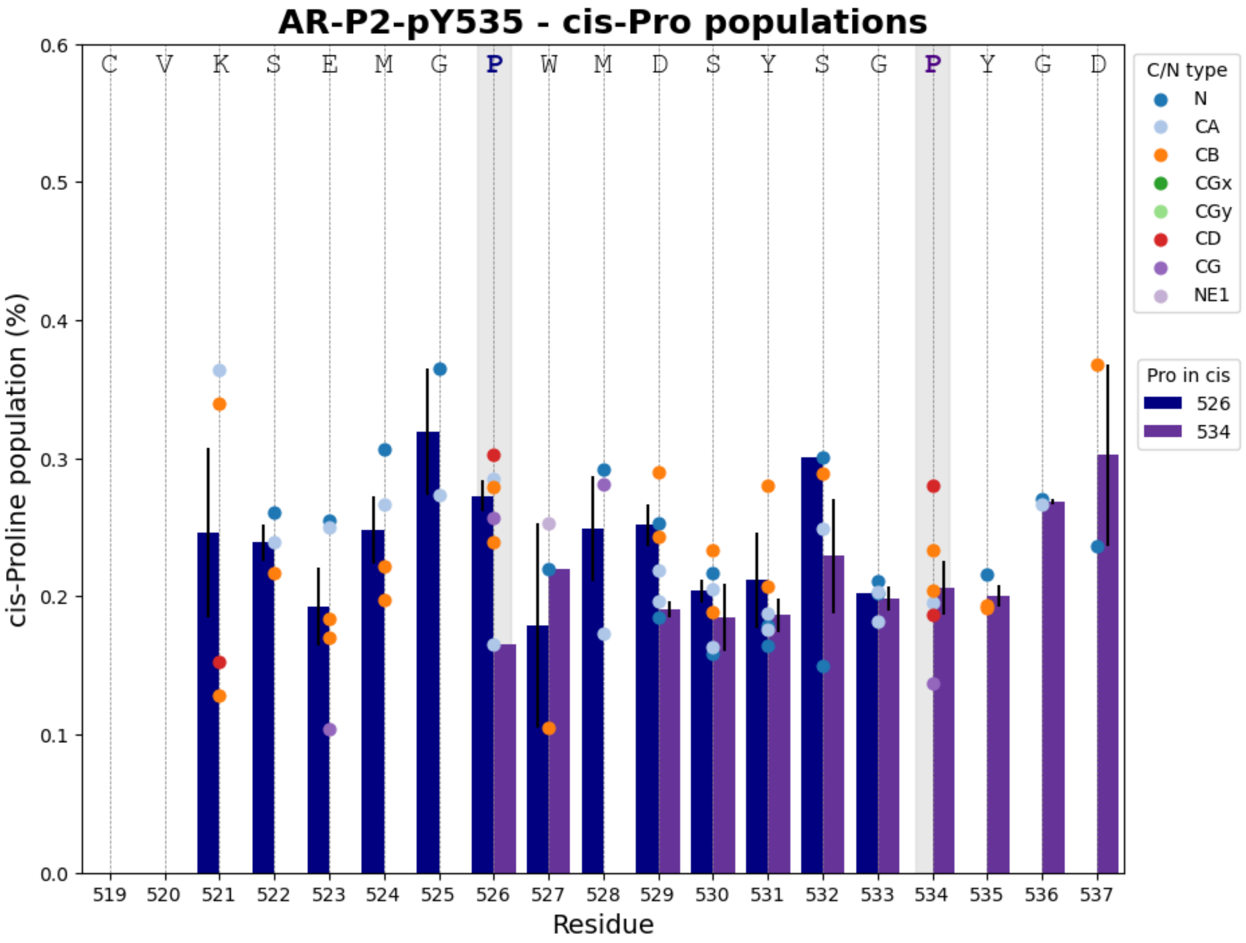


Figure S13: cis-Proline populations for each of the prolines within the P2 peptide arboring the pY535 modification (phosphorylated Y535). The relative populations of trans and cis proline isomers were estimated by comparing the intensities of correlation peaks corresponding to the same nuclei in distinct conformations associated with different proline isomers, using the $^{15}N$-$^{1}H$ and $^{13}C$-$^{1}H$ HSQC spectra. Scatter dots show the contribution of individual nuclei. When different correlation peaks could be used for one residue, bars show the average of all nuclei contribution for a given residue and cis Proline conformation. The uncertainty in the population estimates, for one residue and one proline conformation, was calculated as the standard deviation of all nuclei contribution, corrected using the appropriate Student's t coefficient for a two-sided 95% confidence interval. Uncertainty contributions arising from the signal-to-noise ratio of individual peaks were found to be negligible relative to the variability between population estimates obtained from different correlations and were therefore neglected.
